# Adapting macroecology to microbiology: using occupancy modelling to assess functional profiles across metagenomes

**DOI:** 10.1101/2021.06.21.449349

**Authors:** Angus S. Hilts, Manjot S. Hunjan, Laura A. Hug

**Author notes:** Department of Functional and Evolutionary Ecology, University of Vienna, Vienna, Austria (Angus S. Hilts).

## Abstract

Metagenomic sequencing provides information on the metabolic capacities and taxonomic affiliations for members of a microbial community. When assessing metabolic functions in a community, missing genes in pathways can occur in two ways: the genes may legitimately be missing from the community whose DNA was sequenced, or the genes were missed during shotgun sequencing or failed to assemble, and thus the metabolic capacity of interest is wrongly absent from the sequence data. Here, we borrow and adapt occupancy modelling from macroecology to provide mathematical context to metabolic predictions from metagenomes. We review the five assumptions underlying occupancy modelling through the lens of microbial community sequence data. Using the methane cycle, we apply occupancy modelling to examine the presence and absence of methanogenesis and methanotrophy genes from nearly 10,000 metagenomes spanning global environments. We determine that methanogenesis and methanotrophy are positively correlated across environments, and note that the lack of available standardized metadata for most metagenomes is a significant hindrance to large-scale statistical analyses. We present this adaptation of macroecology’s occupancy modelling to metagenomics as a tool for assessing presence/absence of traits in environmental microbiological surveys. We further initiate a call for stronger metadata standards to accompany metagenome deposition, to enable robust statistical approaches in the future.

## Importance

Metagenomics is maturing rapidly as a field, but is hampered by a lack of available statistical tools. A primary area of uncertainty is around missing genes or functions from a metagenomic datasets. Here, we borrow an established modelling approach from macroecology and adapt it to metagenomic datasets. Rather than multiple sampling trips to a specific area to detect a species of interest (e.g., identifying a cardinal in a forest), we leverage the enormous amount of information within a metagenome and use multiple gene markers for a function of interest (e.g., subunits of an enzyme complex). We applied our adapted occupancy modelling to a case study examining methane cycling capacity. Our models show methanogens and methanotrophs are both more likely to co-occur than be present in the absence of the other guild. The lack of consistent and complete metadata is a significant hurdle for increasing the statistical rigour of metagenomic analyses.

## Background

Environmental microbiology has been a methods-limited field since its conception. Paradigm shifts in our understanding of microbial diversity and the ecological importance of microbes have come hand in hand with new techniques – from the invention of the microscope to high throughput sequencing strategies. Each ground-breaking technique then undergoes improvements, refinements, and matures into a standard approach. Metagenomics, or total community sequencing (1, 2), has demonstrated that the diversity of life on Earth is far greater than was previously believed (3–6). With metagenomics now a standard method of investigation, and public databases filling with deep sequencing datasets from sampling locations around the globe, there is a growing need for statistical tests that can be applied to anchor conclusions based on metagenomic data and to understand patterns from incomplete data (see (7) for a discussion of current statistical methods).

A current challenge is placing the information obtained from metagenomic sequencing studies into a greater ecological context. Microbial roles in geochemical cycles and their contributions to microbial communities can be predicted from the annotated genes within metagenomes. In generating metabolic predictions for a community, a major challenge is interpreting gene absences. Genes may be absent from a genome or metagenome because they are genuinely not encoded by the genome(s) (*i.e.*, true negatives) or because the sequencing depth or assembly failed to capture genes from the community (*i.e.*, false negatives). There are currently no options for robustly modelling which of these gene absence scenarios is most likely.

This question of how to assess ecological relationships is not new. Macroecologists have studied the relationships of organisms and populations for over a century. The macroecology field has developed a wealth of statistical models, ranging from simple to complex (the founding of the journal Ecological Modelling in 1975 is a testament to this focus). In addition, since the 1960’s there has been a movement in macroecology to develop and enforce standards for metadata collection and quality assurance (8). As a result, the foundation for ecological modelling is in place, but microbial ecologists must now adapt these ideas to their own datasets.

Our research sought to adapt occupancy modelling to metagenomic data. The occupancy model was developed by Mackenzie *et al.* (9) to address an important question for any detection-based study: how can missed detections be accounted for? The model addresses the issue that a non-detection could mean that the subject of interest was not present (*i.e.*, a true negative) or that the observer failed to detect it (*i.e.*, a false negative). The underlying idea of occupancy models is that detection can be modelled as two statistical processes: the probability of the species of interest being present at the given site and, given that it is present, the probability that it was observed (9). Two parameters are used to model these, the proportion of sites occupied (denoted Ψ) and the probability of detection given the species is present at the site (denoted *p*). The value of Ψ can be estimated by counting the number of sites at which the species occurs and dividing it by the total number of sites visited. The value of *p* can be estimated by revisiting a site at which the species is known to be present multiple times, and dividing the number of times the species was observed by the total number of visits. A theoretically perfect model would have unique detection and occupancy parameters for every site, but this model would be extremely challenging to calculate, lack flexibility, and have little, if any, predictive power. Instead, the parameters Ψ and *p* can be generalized (*i.e.*, can be assumed to be equal for all sites and surveys), if the following five assumptions are met:

i. The closure assumption, which states that there is no chance of the occupancy state changing between sampling occasions for the site, within the same season
ii. The probability for occupancy is the same across all sites or is otherwise modelled with appropriate covariates
iii. The probability for detection is the same across all sites or is otherwise modelled with appropriate covariates
iv. The detection at each site is independent of detection at other sites
v. There are no false positives

These assumptions must be systematically considered when reinterpreting the model for sequence-based datasets.

The original occupancy model was designed to assess the occupancy of a single species, but its applications have since been expanded. Multi-species models allow exploration of other species of interest (10) with more recent models allowing multi-species modelling without assuming a dominant partner (11).

Here, we apply occupancy models to metagenomic datasets. We develop a method for deriving replicate sampling for functions of interest from a single metagenome, and assess each of the five model assumptions under metagenomic data. We applied this approach as a proof-of-principle to the global methane cycle, using a dataset of 9,629 metagenomes spanning global environments. From these metagenomes, we identified and curated markers for methanogenesis (McrABG) and methanotrophy (PmoABC) as proxies for predicting these functions in an environment, and assessed cooccurrence patterns for these two critical components of the methane cycle.

## Methods

### Reference proteins retrieval

Reference sequences for the proteins of interest (methanogenesis: McrA, McrB, McrG; methanotrophy: PmoA, PmoB, PmoC) as well as known homologs with alternate functions (AmoA, AmoB, AmoC) were retrieved from the Genome Taxonomy Database (GTDB; 12) using AnnoTree (13; Accessed February 11^th^, 2019) and the Kyoto Encyclopedia of Genes and Genomes Orthology (KO) based on KO numbers (Table 1, 14). Data were downloaded in CSV format. Any sequences that were metagenome-derived or for which taxonomy was not resolved to the species level were removed, unless there was supporting literature for the protein as a true representative of the function of interest. Sequences were imported into Geneious (v. 11.0.2; 15), aligned with MUSCLE v. 3.8.425 (16), and maximum likelihood trees inferred using FastTree v. 2.1.5 (17) to assess the quality of the reference sets. Final reference datasets’ accession numbers and host names are listed in Supplemental Table S1.

**Table 1:** Protein sequences identified for reference sets (all) as well as from annotated metagenomes (MCR and pMMO complexes only). Note that Pmo and Amo proteins share KO identifiers. Asterisks signify the active site-containing subunit of the complexes.

| Protein symbol | Gene name | KO identifier |
| --- | --- | --- |
| McrA* | methyl-coenzyme M reductase alpha subunit | K00399 |
| McrB | methyl-coenzyme M reductase beta subunit | K00401 |
| McrG | methyl-coenzyme M reductase gamma subunit | K00402 |
| PmoA | particulate methane/ammonia monooxygenase subunit A | K10944 |
| PmoB* | particulate methane/ammonia monooxygenase subunit B | K10945 |
| PmoC | particulate methane/ammonia monooxygenase subunit C | K10946 |
| AmoA | Ammonia monooxygenase subunit A | K10944 |
| AmoB* | Ammonia monooxygenase subunit B | K10945 |
| AmoC | Ammonia monooxygenase subunit C | K10946 |

### Metagenome-derived gene collection and curation

Data was retrieved from the Joint Genome Institute’s portal for Integrated Microbial Genomes and Microbiomes (JGI IMG/M; accessed between December 14^th^, 2018 and February 22^nd^, 2019). All sequences annotated with KOs of interest (Table 1, top) were downloaded in FASTA amino acid format along with metadata tables for the metagenomes from which they were derived. Sequences retrieved from IMG/M were first filtered based on size in a protein-specific manner, with passing sequences falling between a minimum length of the smallest sequence in the reference set less 50 amino acids and a maximum length of 50 amino acids longer than the longest reference sequence for that protein. Sequences were then screened for known homologs with off-target functions (*e.g.*, butyrate-active Mcr homologs, ammonia monooxygenases).

References sequences were labelled as either true or false positives and then searched against a local copy of the UniRef50 database (release 2019_04) using DIAMOND BLASTp v0.9.24 (18). Any sequences within the Uniref50 database matching to either a true or false positive were removed. The labelled true and false positive reference sequences were then included in this modified database. The length-curated sequences from IMG/M were searched against the modified database using DIAMOND BLASTp. Sequences for which the top hit was not one of the labelled true positives were removed from further analyses. The remaining sequences from IMG/M were then imported into Geneious, aligned to the reference sequences using MUSCLE, and maximum likelihood trees were inferred with FastTree from the alignments. From these trees, sequences which did not cluster with the reference sequences, were excessively divergent, or were on extremely long branches, were removed.

### Occupancy table construction

A table containing a row for each metagenome was generated, with columns for each of the six proteins of interest. Each metagenome for which at least one sequence of a given protein remained after curation were marked with a ‘1’ in the appropriate column, otherwise they were marked with ‘0’. Here, ‘1’ represents a detection and ‘0’ represents a non-detection, regardless of total number of detections for a given protein within a given metagenome. These were collectively referred to as the detection histories for each site (Table S2). This table was imported into R (v3.5.3) along with a table containing metadata for each metagenome, including geographic coordinates (longitude and latitude), ecosystem type (coded as one of ‘host-associated’, ‘environmental’, or ‘engineered’), and the date that the samples were uploaded to IMG/M, converted into days since 2006-01-01. These variables were chosen for their completeness on IMG/M. The data were aggregated into three different datasets. The first dataset had no aggregation, with each metagenome treated as a separate sample/site. The second dataset aggregated metagenomes with identical geographic coordinates into a single site, where any metagenome encoding a gene of interest was sufficient to code that gene as a 1/presence for the given aggregated site. The third dataset aggregated metagenomes with identical geographic coordinates and ecosystem types into a single site, under the assumption that a shift from environmental/engineered/host-associated implied a physical separation of samples despite shared geocoordinates (*e.g.*, a sample from a cow rumen and from soil in the same pasture are not likely to be directly impacting one another, compared to samples from different depths in a soil core). This aggregation was achieved using the ‘dplyr’ (v0.8.0.1) package (19). Finally, sites with missing metadata were removed, to allow comparison of sites.

### Single-species occupancy modelling

Single-species occupancy models were used as a preliminary analysis. The “species” were defined as the functions of interest (methanogenesis and methanotrophy). Surveys (*i.e.*, the repeated sampling events) were defined as the individual genes encoding the enzymes of interest. The R package ‘unmarked’ (v0.12-3) (20) was used for occupancy modelling. There were six model sets in total, with a model for each function and each of the aggregated data sets described above. To run the models, data were loaded into ‘unmarkedOccuFrame’ objects, which combined the detection history data (*i.e.*, the presence/absence table, Table S2) with the site-level metadata. The function ‘occu’ was used with default parameters, other than the engine parameter, which was set to ‘C’. Models with different statistical parameters were run and compared using the Akaike Information Criterion (AIC) (21). Where applicable, the occupancy data were plotted against different covariates with 95% confidence intervals.

### Multi-species occupancy modelling

Multi-species occupancy models were fit according to the Rota *et al.* (11) model using the function ‘occuMulti’ in the ‘unmarked’ package for R in a manner similar to the single-species modelling. Data for both metabolic functions were combined into an ‘unmarkedOccuFrameMulti’ object. Models with different parameterizations (Table 2) were run and compared using AIC.

**Table 2:** Occupancy model parameters for MCR and pMMO datasets. Separate models were developed for each aggregated set of global metagenomes under each of the possible Ψ values, with p held to 1 or as the square root of the metagenome’s add date (as a count from 2006-01-01). Results are ordered by ascending AIC for MCR. AIC = Akaike Information Criterion

| Model | #<br>p | MCR |  | pMMO |  |
| --- | --- | --- | --- | --- | --- |
| | | AIC | ΔAIC | AIC | ΔAIC |
| **Metagenomes as sites (9,420)** |  |  |  |  |  |
| p ~ 1, $\Psi$ ~ Ecosystem | 4 | 9,691.23 | 0.00 | 15,669.51 | 0.00 |
| p ~ sqrt(Numeric.Add.Date), $\Psi$ ~ Latitude | 4 | 9,934.10 | 242.87 | 15,678.85 | 9.35 |
| p ~ 1, $\Psi$ ~ Latitude | 3 | 9,958.31 | 267.08 | 15,929.51 | 260.00 |
| p ~ sqrt(Numeric.Add.Date), $\Psi$ ~ 1 | 3 | 9,982.88 | 291.65 | 15,807.38 | 137.87 |
| p ~ 1, $\Psi$ ~ 1 | 2 | 10,007.15 | 315.91 | 16,053.68 | 384.17 |
| **Aggregated by geocoordinates and environment (1,229 sites)** |  |  |  |  |  |
| p ~ 1, $\Psi$ ~ Ecosystem | 4 | 1,555.15 | 0.00 | 2,284.47 | 0.00 |
| p ~ 1, $\Psi$ ~ Latitude | 3 | 1,594.58 | 39.43 | 2,329.47 | 45.00 |
| p ~ sqrt(Numeric.Add.Date), $\Psi$ ~ Latitude | 4 | 1,596.54 | 41.39 | 2,303.10 | 18.63 |
| p ~ 1, $\Psi$ ~ 1 | 2 | 1,619.65 | 64.50 | 2,330.83 | 46.36 |
| p ~ sqrt(Numeric.Add.Date), $\Psi$ ~ 1 | 3 | 1,621.61 | 66.46 | 2,305.06 | 20.58 |
| **Aggregated by geocoordinates only (1,202 sites)** |  |  |  |  |  |
| p ~ 1, $\Psi$ ~ Latitude | 3 | 1,578.66 | 0.00 | 2,297.88 | 0.00 |
| p ~ 1, $\Psi$ ~ 1 | 2 | 1,605.03 | 26.37 | 2,299.55 | 1.67 |

## Results and Discussion

### Occupancy modelling as applied to metagenomes

In adapting occupancy modelling to metagenomics, the question for microbial ecologists becomes one of identifying parallels between macroecological and microbiological datasets or systems. There are several key aspects that need to be considered here, and in each of these aspects, the solutions may not be universal or applicable to all studies. First, the definition of a site needs to be considered. In addition to this, how a sample unit is defined is important, particularly when considering the cost and labour involved in re-sampling a metagenome. Finally, the applicability of the assumptions underlying occupancy modelling must be determined.

#### Site definition

There are multiple possible definitions of a site, with the simplest being defining a single sample taken for metagenomics as a site. This has the advantage of unambiguously delineating the site, but also poses some disadvantages. For example, a hypothesis exploring cooccurrence of two or more species may not require that the species be present in exactly the same sample, but that they be present within the same system (*e.g.*, a soil core, lake depth profile, or host gastrointestinal system). In these cases, it may be better to define a geographic location or geographic feature as a single site (*e.g.*, the same forest, lake, host), aggregating all associated metagenomes for the defined site. Ultimately, the definition of a site will be dependent on the specific questions of each study, but a clear definition is an important pre-analysis requirement.

#### Sample unit

Another key component of occupancy modelling is that re-sampling a site yields an estimation of the observer’s ability to detect the subject of interest (the variable *p* within the models). Thus, a definition of a re-sampling event, or replicate of a surveyed site must be established. When working with metagenomes, there are several issues with re-sampling. It is expensive, labour-intensive, and time-consuming to obtain samples, isolate DNA, and perform sequencing analyses. Furthermore, it is not always clear what might constitute a replicate sample. Microbial communities mere centimetres apart can be substantially and meaningfully different (22), and by sampling a site, community composition can be changed due to disturbances. In contrast to macroecological observational surveys, however, each metagenome provides an enormous volume of data (Figure 1), containing information from the genomic sequences of hundreds or thousands of microbial populations (23, 24). Given the depth of information available, we instead propose re-sampling a single metagenome for multiple genetic markers of a function or group of interest. Each independent marker is then considered a replicate survey for the target function or taxonomic group.

**Figure 1:**
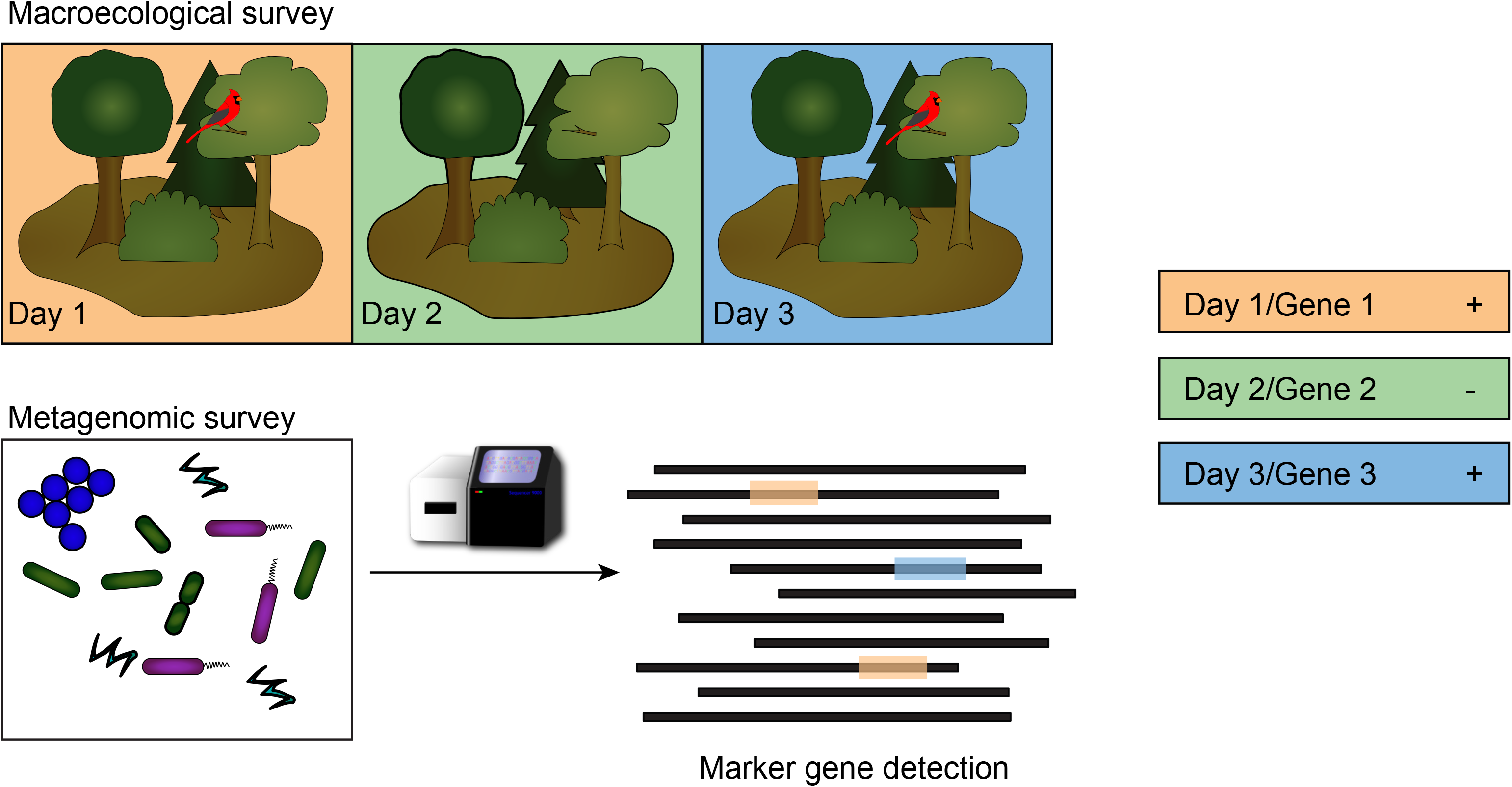
Schematic depicting connections between occupancy sampling for macroecologists compared to the proposed method for metagenomic datasets. Where macroecological occupancy sampling requires repeated observation (days 1, 2, 3) of the same environment (forest) for presence/absence of a species of interest (bird), we propose using molecular markers (genes 1, 2, 3) for an activity or lineage of interest that allow multiple observations of presence/absence from within a single metagenomic dataset.

From this, a critical step in the application of occupancy models to microbial ecology becomes the decision of which marker genes to use. Genes must be universally and uniquely associated with the target function or taxonomic group. For a target function, an obvious choice are genes encoding different, required subunits of an enzyme complex associated with the function of interest. For a taxonomic group, core genes specifically and universally associated with the group of interest would be required. Occupancy models are explicitly designed to avoid the problem of false-negatives, but as a result they are biased toward false-positives. As a result, it is important that there are no, or very few, false positives in the underlying datasets, so whether a marker can be separated into true or false positives should be a first consideration. How false positives are filtered out will depend on the selected marker genes, but considerations such as active site, conserved residues, secondary and tertiary structure modelling, length requirements, homology to characterized proteins, and phylogenetic placement can be applied. There are a growing number of applications of occupancy modelling incorporating false positives into the modelling process for macroecological studies (25, 26), but at base, curation of potential false positives from the data is an important step. The use of multiple marker genes as “re-samplings” provides the advantage that multiple metagenomic samples are not needed, and also means that several of the assumptions of occupancy models are met “for free” (see next).

#### Occupancy modelling assumptions

Occupancy modelling separates successful detections into two processes: first, the species of interest must be present at the survey site, and second, given that it is present, it must be successfully detected. These processes are modelled as two probabilities, denoted Ψ and *p* (9). The first parameter, Ψ, represents the proportion of occupied sites, while the latter, *p*, represents the probability that, given the species is present, it is successfully detected. A theoretically perfect model could have a unique detection and occupancy parameter for each site, but this would sacrifice flexibility. Instead, the model can be generalized using certain assumptions, which restrict these parameters to single values across all sites. Mackenzie *et al*. (9) proposed five assumptions that, if met, allow for this generalization to be applied. In brief, each assumption seems reasonable to apply to metagenomic datasets, but there are specific points that require consideration or definition.

##### i) The closure assumption, which states that there is no chance of the occupancy state changing between sampling occasions for the site, within the same season

The first assumption is met without ambiguity when using multiple genetic markers from a single metagenome as the sampling units. Detection of these markers is conducted on the same metagenome, from the same extracted DNA, taken at the exact same time. This is an advantage of metagenomic datasets compared to macroecological datasets, where the observer must return to a site over multiple separate occasions.

##### ii) The probability for occupancy is the same across all sites, or is otherwise modelled with appropriate covariates and iii)The probability for detection is the same across all sites, or is otherwise modelled with appropriate covariates

The second (ii) and third (iii) assumptions are more challenging. It is unlikely that detection and occupancy are uniform across all sites. In particular, the occupancy state is expected to change as a result of different environmental conditions, such as pH, temperature, and organic carbon content of the site. It may be more reasonable to assume that detection probabilities are uniform, but even this is unlikely. Factors affecting detection probabilities include challenges in DNA extraction for difficult matrices or resistant microorganisms as well as variations due to changing sequencing technologies, where greater sequencing depths enable better assemblies and a higher likelihood of detecting a function or taxon at low abundance. For occupancy modelling, variation in detection and occupancy are acceptable, so long as they are modelled with covariates. In this context, a covariate might be metagenome size, pH, total organic carbon, elevation above sea level, or other factors that would influence occupancy and detection for the trait or taxon of interest. Identifying useful covariates was one of the major challenges in our methane cycle case study, described below. Metadata reported for metagenomes was sporadic and incomplete, meaning that there were very few variables that could be used as covariates. Better data deposition standards have the potential to rapidly remedy this problem.

##### iv The detection at each site is independent of detection at other sites

Assumption (iv) is a reasonable assumption for metagenomic data. Unless samples were cross-contaminated, metagenomes should have no impact on the content of other metagenomes and thus can be treated as independent from one another. This holds true if aggregating metagenomes into larger scale samples (*e.g.*, a soil core), as long as each set of metagenomes are from samples/sites that do not meaningfully interact.

##### v) There are no false positives

This assumption also poses a challenge. The best solution is to employ procedures that carefully curate the detections prior to modelling, as described above. Searching against established databases, the use of annotation pipelines, multiple sequence alignments, and phylogenetics can be used to minimize the number of false positives. Further case-specific information can be valuable, such as identifying key conserved residues at active sites, or motifs thought to be conserved across all members of the group of interest. By incorporating this sort of information and applying rigorous filtering to the data, false positives can be minimized or eliminated.

### Occupancy modelling and the global methane cycle

We took our theorized definitions of sites, sampling units, and the assumptions required for occupancy modelling and applied them to a case study examining the cooccurrence of methanogenesis and methanotrophy across global environments.

Two microbial groups are the main controls on biological methane cycling and the flux of methane emissions. The majority of methane emissions originate from methanogenic archaea (Schimel, 2004). The other group of organisms implicated in the methane cycle are the methane oxidizers. Methane oxidizing organisms largely fall into two categories: the bacterial methanotrophs (Hanson and Hanson, 1996), and the archaeal anaerobic oxidizers of methane (AOM; Hinrichs et al., 1999; Cui et al., 2015). Understanding the cooccurrence patterns of methanogens and methanotrophs across global environments would provide insight into methane emission fluxes and spotlight regions where microbial control of methane emissions is tipped toward higher emissions (*i.e.*, sites with methanogens and no associated methanotrophs). Using 9,629 publicly available metagenomes from a wide variety of global environments, we applied occupancy modelling to metagenomic data to strengthen cooccurrence analyses of methanogens and methanotrophs.

In defining a “site” for the occupancy models, we tested three approaches. First, we tested the naive approach of each metagenome equating to a single site. However, methanogens and methanotrophs differ in their oxygen requirements: methanogens are obligately anaerobic whereas methanotrophs are generally understood to be microaerophilic (27, 28), though this paradigm is shifting with increased reports of methanotrophy in anoxic conditions (29, 30). Given this difference, we would not generally expect methanogens and methanotrophs to occur at the exact same location, but their presence within the same system would be informative. To address this, our second approach was to aggregate metagenomes from the same geographic coordinates (latitude and longitude) as a single site, meaning soil cores and depth profiles were merged. Our third and final approach was to separate the aggregated metagenomes in the second approach if their environmental coding differed between environmental, engineered, and host-associated.

For sampling units, we required a set of genes that could act as proxies for function and be used to emulate re-sampling an environment, within a single metagenome. We selected subunits of the catalytic enzyme complexes responsible for the final step of methanogenesis and the first step of methane oxidation: methyl- coenzyme M reductase (MCR; subunits McrABG) and particulate methane monooxygenase (pMMO; subunits PmoABC). Both enzyme complexes are built from three distinct and necessary subunits. All known methanogens contain *mcr* genes in their genomes (31, 32), which encode the MCR complex responsible for the final step in methane formation (33, 34). The *mcr* operon contains a suite of genes, but the three that encode for the MCR complex are *mcrA*, *mcrB*, and *mcrG*, which encode the α, β, and γ subunits, respectively (35). The MCR enzyme complex is an α_2_β_2_*γ*_2_ hexamer (36). The *mcrABG* genes are frequently used in combination as methanogenic markers (37–39) and can be used in place of 16S rRNA genes to infer phylogeny of methanogens (40). The oxidation of methane is catalyzed by methane monooxygenases (MMO), which have two forms, soluble and particulate (41). The soluble form (sMMO) is localized to the cytoplasm, while the particulate form (pMMO) is membrane bound (42). We chose the pMMO enzyme over sMMO as a methanotrophic marker because the soluble form usually occurs alongside the particulate form, but the reverse is not necessarily true. The pMMO complex is encoded by the *pmoCAB* operon (43, 44). These genes, especially *pmoA*, are used as functional markers for aerobic methanotrophy (42, 45, 46). While the *pmo* operon is present in most methanotrophs, it has recently been found to be absent in some species (47, 48; for a review see (46)), meaning this marker may lead to false negatives. Another source of false negatives in our analyses is anaerobic oxidation of methane by the ANME archaea (49, 50). The ANME archaea use the methanogenic pathway in reverse, including McrABG, to catalyze the oxidation of methane (51–53). These methane-oxidizing archaea are fascinating and likely important players in methane cycling (54), but there is growing evidence the ANME can also produce methane (55), meaning their McrABG are ambiguous markers for methane cycling, and we wanted a straightforward initial case study. As a result, for the work described below our focus was on oxidation of methane performed by bacteria. The main relevance of the ANME archaea for this research was that any McrABG proteins that were closely associated with the ANME MCR were removed from the methanogenesis datasets.

Excluding sMMO and ANME MCR from our methanotrophic survey means we have not captured the total methane oxidation capacity from the surveyed environments, but for this proof of principle exercise, the advantages of having an equal number of marker genes and a single set of markers per function of interest outweighed the potential for false negatives.

Reviewing the 5 assumptions to allow for generalized Ψ and *p* parameters in our occupancy model, assumption (i) is met, as each metagenome or aggregated set of metagenomes was static in time and searched simultaneously for the six marker genes. Assumptions (ii) and (iii) are not met with the raw data, and so we used covariates to model factors impacting occupancy (ii) and detection (iii) probabilities. Assumption (iv) is met, as detections between sites are independent, given the reasonable assumption that each metagenome stemmed from independent DNA extractions from isolated samples (*e.g.*, sample handling minimized cross-contamination). Assumption (v), that there are no false positives, required significant dataset curation of our candidate marker genes.

The data set used consisted of 9,629 metagenomes from across every continent on Earth (Figure 2). The metagenomes were classified into three broad categories: 80.6% were from environmental samples, 13.3% were host-associated samples, and 6.2% were from engineered environments. From these datasets 261,869 sequences were identified as candidate matches to the target genes (based on KEGG KO annotations). After size-filtering sequences size to remove partial and truncated sequences, 59,390 candidate sequences remained. Filtering using DIAMOND BLASTp further reduced the set, leaving 29,308 sequences. Finally, after manual assessment for phylogenetic congruence, 27,730 of the initial sequences remained. This curation was important to minimize false positives to satisfy assumption (v). Following curation, the frequencies of each complex’s subunits were similar within different environmental categories. Single species models were run for each of the two functions (methanogenesis and methanotrophy) (Table 2). In nearly all cases, the null model (*i.e.*, models run without any covariates) explained the least variability in the data. The available covariates were ecosystem type, latitude, and date of metagenome deposition. Where applicable, ecosystem type as a covariate for the occupancy state contributed to the best model (Table 2). In addition to this, the latitude seemed to offer some small improvement. The square-root-transformed add date, counted as days since 2006-01-01, improved models when used as a covariate for the detection probability (Table 2).

**Figure 2:**
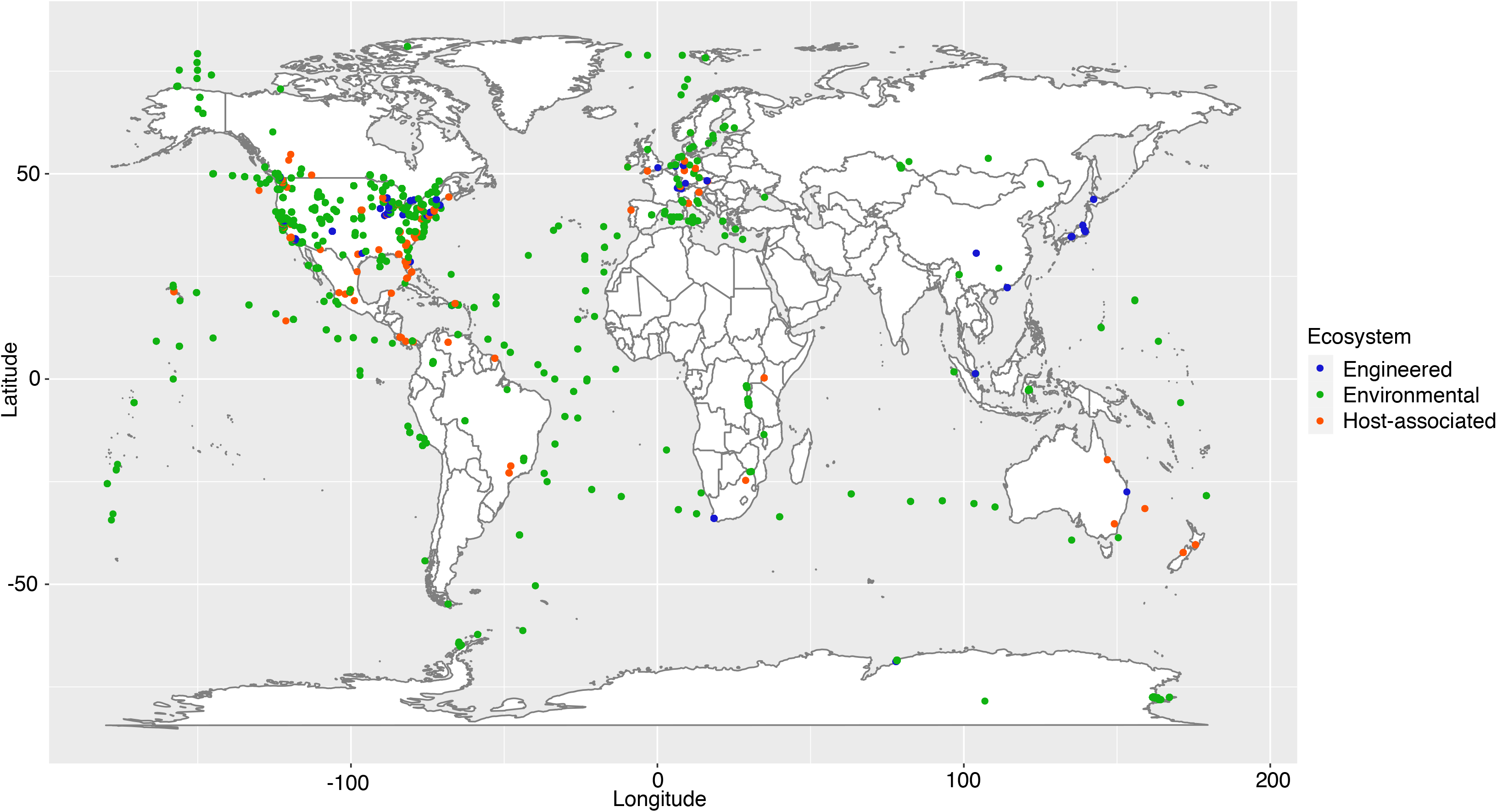
Geocoordinates associated with all metagenomes examined for methanogenesis and methanotrophy marker enzymes. All continents were included within this survey, but there is a clear sampling bias towards North America and Europe.

In general, the models predicted that methanogens occupied a larger portion of engineered sites than methanotrophs. In contrast, a higher proportion of environmental sites were predicted to be occupied by methanotrophs than methanogens, and the proportion of host-associated sites occupied by each group was similar (Figure 3). For both functional groups, the proportion of sites occupied appeared to increase with latitude. Interestingly, this trend followed from extreme Southern latitudes through to the extreme Northern latitudes, rather than increasing from the equator toward both poles (Figure S1). This trend may reflect a strong geographic bias in sampling. The majority of available metagenomes were from samples taken from locations in North America and Western Europe (Figure 2), which cover a similar range of latitudes. In practice, this may impact the model estimates of occupancy, biasing them toward detection in better-covered regions of the map. Finally, the estimated detection probability increased with the square-root-transformed dates (Figure S2). This may not mean there was an increase in prevalence of these metabolisms over time, but rather that this covariate captures underlying differences in the data that would be better or more precisely captured by other covariates that were not accessible to us due to incomplete metadata for the submitted metagenomes. For example, the observed increase may be tied to advances in technology, and/or be due to increasing metagenome sizes over time leading to higher detection probabilities of lower abundance functions.

**Figure 3:**
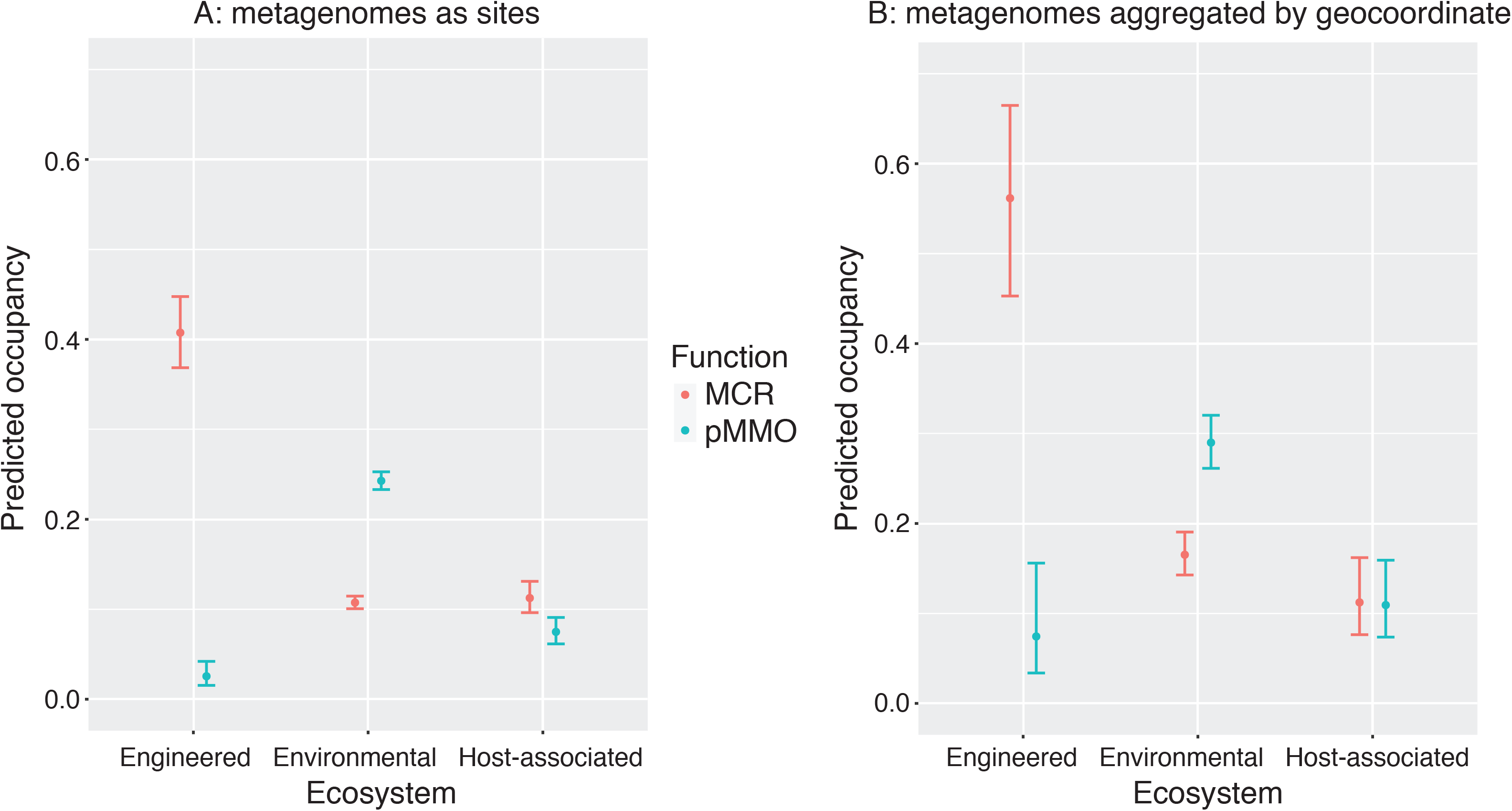
Predicted proportion of sites occupied for both MCR and pMMO. Panel A represents the unaggregated metagenomes while panel B represents aggregated metagenomes where the geocoordinate and ecosytem type were the same. The model used for both plots was *p ~ 1, Ѱ ~ Ecosystem*. Error bars indicate 95% confidence intervals for the estimated occupancies.

In the case of the multi-species models, the ecosystem covariate was found to have the strongest influence in explaining the data, regardless of the dataset used (Table 3). In all cases, the presence of the other functional group in the model increased the estimated proportion of sites occupied for the other group (Figure 4, Figure S3). For the unaggregated data, the occupancy in MCR increased if pMMO was present for the entire range of latitudes, in all three environment types (Figure 4). According to the models, this prediction was most robust for engineered sites. The confidence intervals for both environmental and host-associated sites were much broader, but the same trends were observed. The same trend was observed for pMMO, which increased in occupancy if MCR was present. However, the occupancy of pMMO in engineered sites was very low regardless of the occupancy of MCR. The aggregated data sets were similar in pattern compared to the unaggregated data. However, the confidence intervals were much larger, most notably for the occupancy of pMMO at environmental sites (Figure S3). The data that were aggregated solely by geocoordinates showed the clearest trend where the occupancy of each function increased when the other function was present (Figure S4). A hypothesis of this work was that methanotrophs would be more likely to occur at sites containing methanogens, or would occur at close proximity to sites containing methanogens. The opposite case, where a methanogen would be more likely to occur if a methanotroph was present, is not anchored in our understanding of the biology of these organisms. This makes our model results interesting, since both scenarios are predicted to be true. It may be that external variables controlling the presence or absence of each group are shared, and so the two functional groups coincide because of limitations to their distributions. The size of the confidence intervals across all trials prevent strong conclusions from being drawn. This is not discouraging, as the ability to associate confidence intervals with metagenomic predictions, and to temper conclusions based on statistical analyses is a necessary and important addition to the maturation of metagenomics as a field.

**Table 3.**
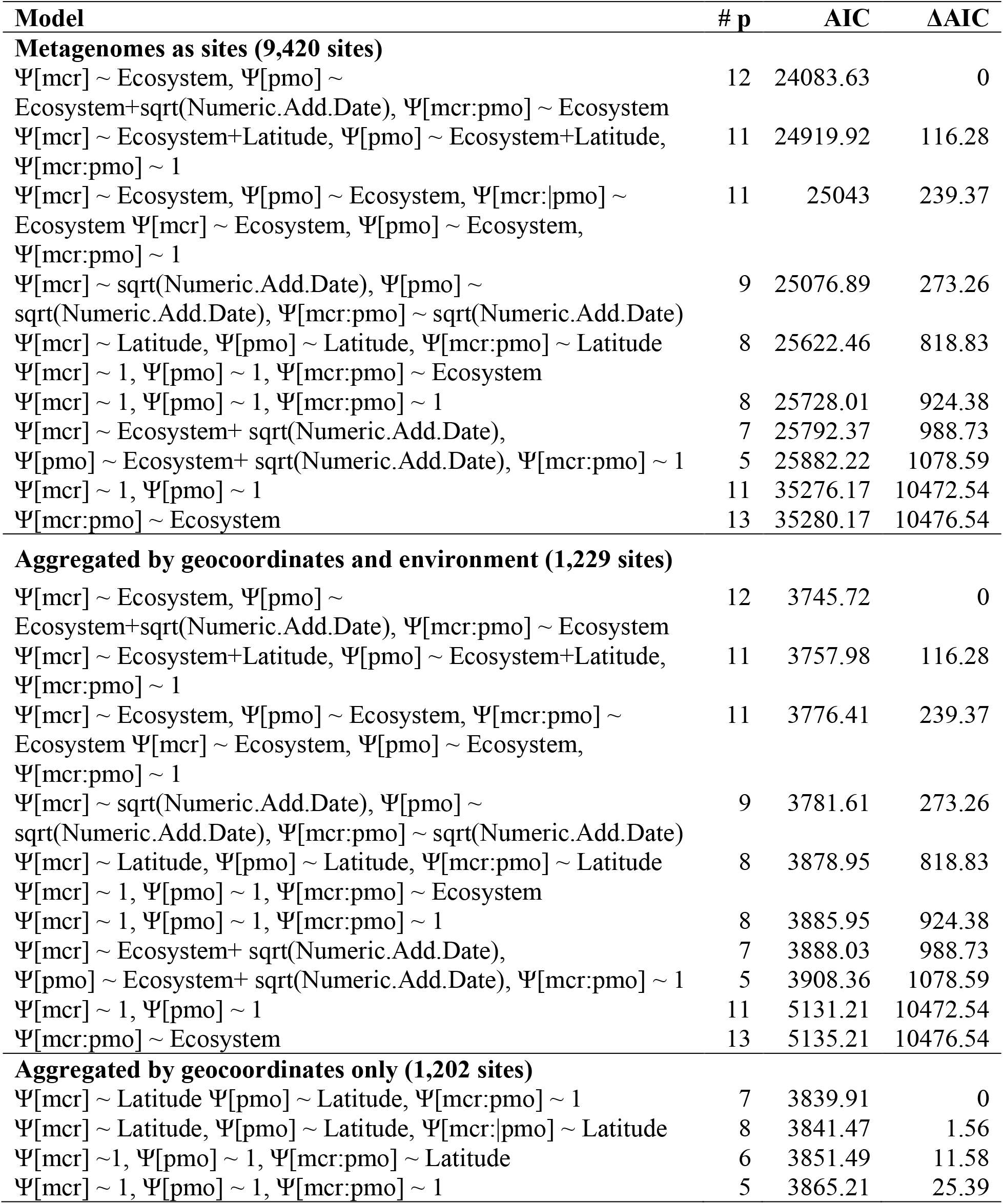
Multi-species occupancy models for MCR and pMMO across global metagenomes. Separate models were developed for each aggregated set of global metagenomes under each of the possible Ψ values, with p held to 1. AIC = Akaike Information Criterion

**Figure 4:**
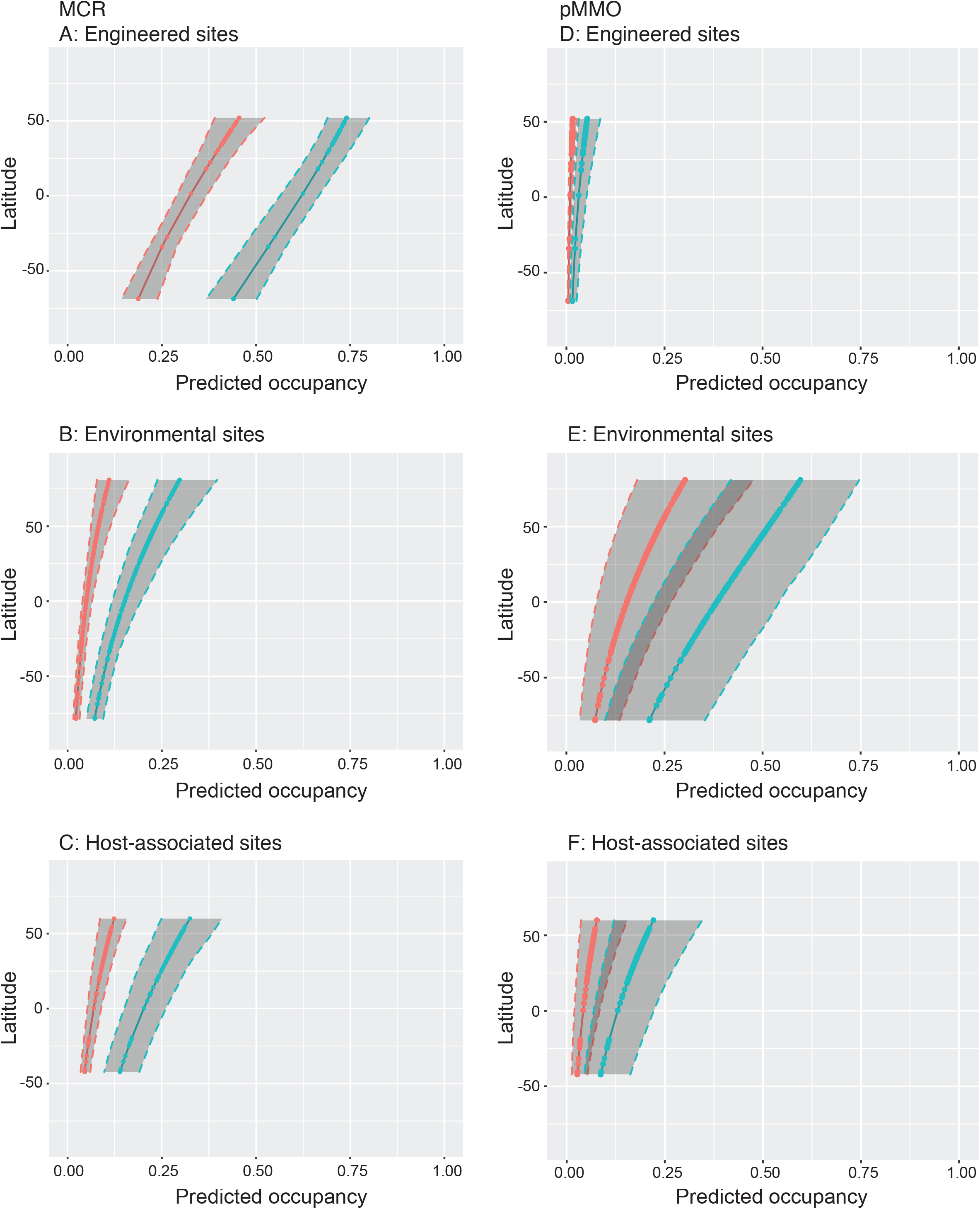
Predicted occupancy for functions of interest given the presence (blue) or absence (red) of the other function. Shaded regions represent 95% confidence intervals for the estimates (solid lines). All sites are unaggregated metagenomes. Panels A-C show the estimated occupancy for MCR, while panels D-F show that of pMMO.

### Metagenomic metadata is a limiting factor for statistical analyses

While our models predicted interesting cooccurrence patterns, we were constrained by the lack of metadata available for the metagenomes of interest. The accuracy of occupancy modeling is dependent on covariates to address changes in detectability and occupancy between sites, particularly continuous covariates that can be used to assess how these values vary as some external parameter does (9). The metadata associated with the metagenomic datasets used here included very few numerical covariates, which were often incomplete across all metagenomes. This seriously restricted our ability to apply these covariates to strengthen the predictive models.

Our models showed occupancy trends associated with the available numerical covariates. We think some of the observed trends may be driven by sampling biases, particularly for the latitude of the sample location. Having more covariates would help to de-conflate the underlying confounding factors that are driving these biases or allow choice of covariates with lower bias levels.

We believe that the remedy to the current lack of covariates is to require better metadata deposition standards. This would not require new or standardized sampling protocols, which would be near impossible to implement across environments. Instead, database administrators could make deposition of more metadata mandatory, to be uploaded alongside sequence datasets. Established and enforced metadata reporting requirements would greatly strengthen modelling practices. This will require defining specific data that must be collected alongside a metagenomic sample at the time of capture. The types of data to include, for example, could be pH, mineral content, oxygen content, and moisture content, among others. Further to this, when categorizing environments in hierarchical classification structures like that of IMG/M, standard definitions need to be established and enforced. This would serve two purposes: first, it would reduce the number of categories employed, allowing use of categorical covariates in statistical models without over-parameterization, and second, it would increase confidence in conclusions, knowing that the categories to which the metagenomes were assigned were well defined without redundancies or overlaps.

## Conclusions

Perfect detection is unlikely to ever be fully achieved in field ecology. The need to properly account for false negatives in detection surveys motivated the development of the occupancy model by Mackenzie, *et al.* in 2002. The method outlined here provides a novel approach for microbial ecologists to apply macroecological occupancy models to microbial metagenomic datasets and to assess microbial interactions on a global scale. The advantage to this approach is that it enables researchers to study a hypothesis of interest using statistical replicates, without the need for expensive re-sampling. We successfully applied occupancy modeling to metagenomic datasets, generating predictions about the cooccurrence patterns of methanogenesis and methanotrophy across global environments. None of the covariates available with this data were particularly compelling, and yet each improved either the occupancy or detection probability model fit when applied. It was challenging to draw strong conclusions in the absence of complete metadata for metagenomes. For metagenomic research to continue to evolve statistical rigour, better data-deposition standards are required to ensure that relevant metadata is available to researchers.

## Acknowledgements

We would like to thank Mr. Jared Ellenor and Dr. Heidi Swanson (University of Waterloo) for invaluable discussions on macroecological approaches to occupancy modelling. We acknowledge the work required in sampling, sequencing, and processing metagenomes, and thank all those involved in generation of the publicly available datasets we examined in this study, as well as the JGI IMG platform for management of these data. This research was supported by an NSERC Discovery grant (2016-03686) and Canada Research Chair to LAH.

## Data availability

All sampled metagenomes were publicly available on the Joint Genome Institute’s Integrated Microbial Genomes database. Accession numbers for reference genomes used for positive and negative control sequences are listed in Supplemental Table S1. Accession numbers and accompanying metadata for the metagenomes used are available in Supplemental Table S2.

## Abbreviations

IMG/M: Integrated Microbial Genomes and Microbiomes
JGI: Joint Genome Institute
KEGG: Kyoto Encyclopedia of Genes and Genomes
KO: KEGG Orthology
*mcr*: Methyl-coenzyme M reductase (referring to the gene)
MCR: Methyl coenzyme M reductase (referring to the enzyme)
*pmo*: Particulate methane monooxygenase (referring to the gene)
pMMO: Particulate methane monooxygenase (referring to the enzyme)

---

**Figure S1:** Predicted proportion of sites occupied versus latitude when using metagenomes as individual sites (panel A), metagenomes aggregated by geocoordinate and ecosystem type as sites (panel B), and metagenomes aggregated by geocoordinates only (panel C). Shaded regions represent 95% confidence intervals for the predicted occupancy (solid lines). Note the smaller confidence intervals in panel A, likely due to inflated sampling, since a given geocoordinate may have multiple samples, which is not the case for panels B and C.

**Figure S2:** Detection probability (*p*) versus date for unaggregated data, as predicted by occupancy models. Dates were converted to a numeric format (counted from 01-01-2006) and square-root-transformed; these were used as the only covariate for prediction. Shaded regions indicate 95% confidence intervals for the estimated detection probabilities (solid lines).

**Figure S3:** Predicted occupancy for functions of interest given the presence (blue) or absence (red) of the other function. Shaded regions represent 95% confidence intervals for the estimates (solid lines). All sites are metagenomes aggregated by their respective geocoordinates and ecosystem types. Panels A-C show the estimated occupancy for MCR, while panels D-F show that of pMMO.

**Figure S4:** Predicted occupancy for functions of interest given the presence (blue) or absence (red) of the other function, with all sites as metagenomes aggregated by their respective geocoordinates and with latitude as a covariate. Shaded regions represent 95% confidence intervals for the estimates (solid lines).

